# Do flowers with specialized morphologies produce more nectar and pollen?

**DOI:** 10.1101/2024.10.10.617618

**Authors:** Tamar Keasar, Levona Bodner

## Abstract

**Premise of the study:** Flower morphology influences the wiring of plant-pollinator interaction networks. Flowers with deep corolla tubes and bilateral symmetry have a narrower pollinator range, hence are considered more specialized than shallow radial flowers. Past inter-specific comparisons revealed positive correlations between flower depth and nectar production rates in a few plant communities, suggesting that specialized flowers may allocate more resources into food rewards for pollinators.

**Methods:** To evaluate this hypothesis, we compiled a global dataset of flower morphology vs. nectar sugar production rates (n = 494 plant species) and per-flower pollen counts (n = 164 species). We applied phylogenetically controlled mixed models to examine the effects of symmetry and tube length on floral rewards.

**Key results:** Corolla tube lengths, symmetry type and their interaction significantly predicted nectar production rates, with the larger effect attributed to tube length. Neither tube length nor symmetry predicted pollen number. Both nectar and pollen production were affected by phylogeny in a larger dataset that included 854 nectar-producing and 1040 pollen-producing species. Plant genus explained more of the variation in nectar production, and less of the variation in pollen production, than plant family.

**Conclusions:** Our findings suggest that visual signals associated with specialized flowers honestly advertise nectar production, but not pollen rewards. Other visual and chemical family-specific floral displays potentially advertise pollen availability to pollinators. We propose experiments to test whether nectar foragers indeed use flower depth as a visual signal that guides their choices of nectar sources.

## INTRODUCTION

To manage, conserve, and restore pollination services in natural communities, is it essential to understand how individual plant and pollinator species interact (Maia et al., 2019). Such interactions are often characterized along a continuum of specialization (Waser et al. 1996; Fenster et al., 2004). Highly generalized interactions represent one end of the range. They involve multiple pollinator taxa that visit each plant species, and numerous flower species visited by each pollinator. At the other end, highly specialized interactions involve an obligatory mutualism between a single plant and a single pollinator. Floral morphological structures that constrain visitors’ access often contribute to a plant’s specialization along the continuum. The length of flowers’ corolla tubes and the type of flower symmetry are well-studied examples of such features. Long-tubed flowers are efficiently probed only by a subset of pollinators with matching mouthparts, and therefore such flowers interact with a narrower range of visitors than short-tubed species (Martins and Johnson, 2013; Stang et al., 2006). Similarly, foragers can feed from zygomorphic (bilaterally symmetrical) flowers only from specific angles, while actinomorphic (radially symmetrical) flowers can be approached from any direction. Consequently, bilaterally symmetrical flowers have fewer pollinator partners than flowers with radial symmetry (Yoder et al., 2020).

In addition to having fewer species of visitors, long-tubed flowers produce more nectar than short-tubed flowers. This trend was recorded in inter-specific comparisons across eight species of Ericaceae (Harder and Cruzan, 1990); hundreds of hummingbird-pollinated plants (Ornelas et al., 2007; Tavares et al., 2016); 25 African hawkmoth-pollinated species (Martins and Johnson, 2013); and among 76 Mediterranean bee-pollinated phrygana plants (Petanidou and Smets, 1995). Two main hypotheses were proposed to explain the correlation between floral tube length and nectar production. The *physiological constraints* hypothesis suggests that long-tubed flowers are simply larger than short-tubed flowers and may, therefore, have more metabolic resources to produce nectar and can store larger nectar volumes (Harder and Cruzan, 1990). An alternative hypothesis, stressing *evolutionary processes*, suggests that long-tubed flowers, being more specialized than short-tubed ones, face higher risks of pollinator limitation. This is because long-tongued pollinators can access both long-tubed and short-tubed flowers, potentially causing both types of flowers to compete for the same pollinators (Martins and Johnson, 2013). Long-tubed flowers consequently experience increased selection to produce food rewards for pollinators (Harder and Cruzan, 1990; Martins and Johnson, 2013). Additionally, and relevant to both hypotheses, both tube length and nectar production rates are influenced by the plants’ phylogeny (Ornelas et al., 2007). For example, perennial species of the Lamiaceae family have high nectar production rates in Mediterranean shrublands than plants of other families (Petanidou and Smets, 1995). Such related species cannot be considered independent records, thus statistical analyses of the relationships between flower traits and rewards should test and correct for shared phylogenetic ancestry. Here, we extend the comparative analysis of nectar production to a larger and global database of plant species. We confirm the previously reported link between nectar production and corolla tube lengths, and test whether nectar production also correlates with floral symmetry type. The evolutionary hypothesis predicts higher nectar production in zygomorphic than in actinomorphic flowers because, being more specialized, zygomorphic flowers are more prone to pollinator limitation. If zygomorphic flowers also tend to be larger than actinomorphic flowers (see below), the physiological constraints hypothesis would make the same prediction. To our knowledge, the relationship between symmetry type and nectar production was so far tested only in a single plant community, and no correlation was found (Ortiz et al., 2021).

We also compiled a dataset of pollen production, which we relate to the plant species’ flower depth and symmetry type. Pollen, which comprises the flowers’ male gametes, is partly transported to stigmas of other flowers during pollination, and partly consumed by pollinators. This dual function generates opposing evolutionary predictions regarding pollen production in specialized flowers. Following the hypotheses presented above for nectar production, flower specialization is predicted to increase pollen amounts, because of selection to attract pollinators and/or because of higher resources available to specialized flowers to produce pollen. On the other hand, morphological specialization improves pollination efficiency by increasing pollinator constancy, reducing between-species pollen transfer, and improving the accuracy of pollen placement on the recipient stigmas. These effects were proposed to reduce pollen wastage, relaxing selection for pollen production (Stephens et al., 2024 for zygomorphic flowers). Similar considerations may likewise reduce pollen production in long-tubed flowers. More generally, pollen numbers are predicted to decline when pollen transfer is efficient, and this expectation was supported in a comparative analysis (Harder and Johnson, 2023). We are not aware of tests of the relationship between floral tube depth and pollen production, and provide a first assessment here. Previous comparisons of pollen production between zygomorphic and actinomorphic flowers yielded inconsistent results. Ortiz et al. (2021) did not detect an effect of floral symmetry type, whereas Cunha et al. (2022) found higher pollen production in bilaterally symmetrical flowers.

Insect visitors invest time and energy in learning to efficiently handle flowers with specialized morphologies (Gegear and Laverty, 1995; Muth et al., 2015; Krishna and Keasar, 2019). Bumble bees increase their attempts to forage on specialized flowers when they contain high rewards (Muth et al., 2015), and after gaining previous experience with other flowers that are difficult to handle (Krishna and Keasar, 2021). These findings suggest that pollinators learn to associate display features of specialized flowers with food rewards, and that such learned associations increase foragers’ preference for specialized flowers over generalized flowers. To further explore this idea, we ask here whether tube length and symmetry are honest signals of food rewards that can potentially attract pollinators to specialized flowers.

## MATERIALS AND METHODS

### Data compilation

We compiled three datasets from published sources (Supplementary Material S1) and from our own field surveys. When we had multiple records for a species, we averaged them. The flower traits dataset contains information on corolla tube length (n = 1967 species) and symmetry type (zygomorphic or actinomorphic, n = 1583 species). The nectar dataset list daily sugar production data for 973 species. Published sources either report sugar production (in mg/flower/day) directly, or provide information on the volume and sugar concentration of nectar produced per day. In the latter cases, we multiplied the volume by the concentration to obtain the mass of sugar produced. The pollen dataset contains mean per-flower pollen grain counts for 1495 species.

### Data analysis

We used phylogenetic linear mixed models (PGLMMs, Li et al., 2020) to test the effects of corolla tube length and symmetry type on nectar and pollen production. We used the default phylogeny provided by the R package V.PhyloMaker2, and selected the ‘scenario 3’ option for building the phylogenetic tree. This analysis was applied to 454 species with data on morphology and nectar production, and 164 species with data on morphology and pollen production. We calculated R^2^ likelihood (R2-lik) values to estimate the relative importance of the fixed factors (tube length and symmetry) and of the random factor (phylogeny) in predicting nectar and pollen production (Ives, 2019).

To determine whether the phylogenetic signal in nectar and pollen production is mainly due to variation among plant genera or among families, we analyzed a subset of the data that included the better-represented families in the dataset (families with >20 species for either nectar or pollen production). This analysis included all species with nectar and pollen production information, whether we had information on their morphology or not. This enabled us to analyze 854 species of 17 families for nectar production and 1040 species of 21 families for pollen production. We used generalized linear models with family Gamma and inverse link functions to relate the plant genus, nested within family, to nectar and pollen production. We used the eta-square measure to evaluate the relative importance of genus vs. family in explaining the variation. Pollen production data were log-transformed before analysis.

We used the packages lme4, phyr, rr2, V.PhyloMaker2 of R version 4.2.2 (R core team, 2022) for the analyses.

## RESULTS

### Distributions of morphological traits

65.1% of the species in the dataset had actinomorphic flowers. The distribution of corolla tube lengths was skewed in both actinomorphic and zygomorphic flowers (Figure 1). Actinomorphic flowers had, on average, shorter corolla tubes than zygomorphic flowers (Figure 1, t-test: t_1103.9 =_ 9.2418, P < 0.0001).

**Figure 1:**
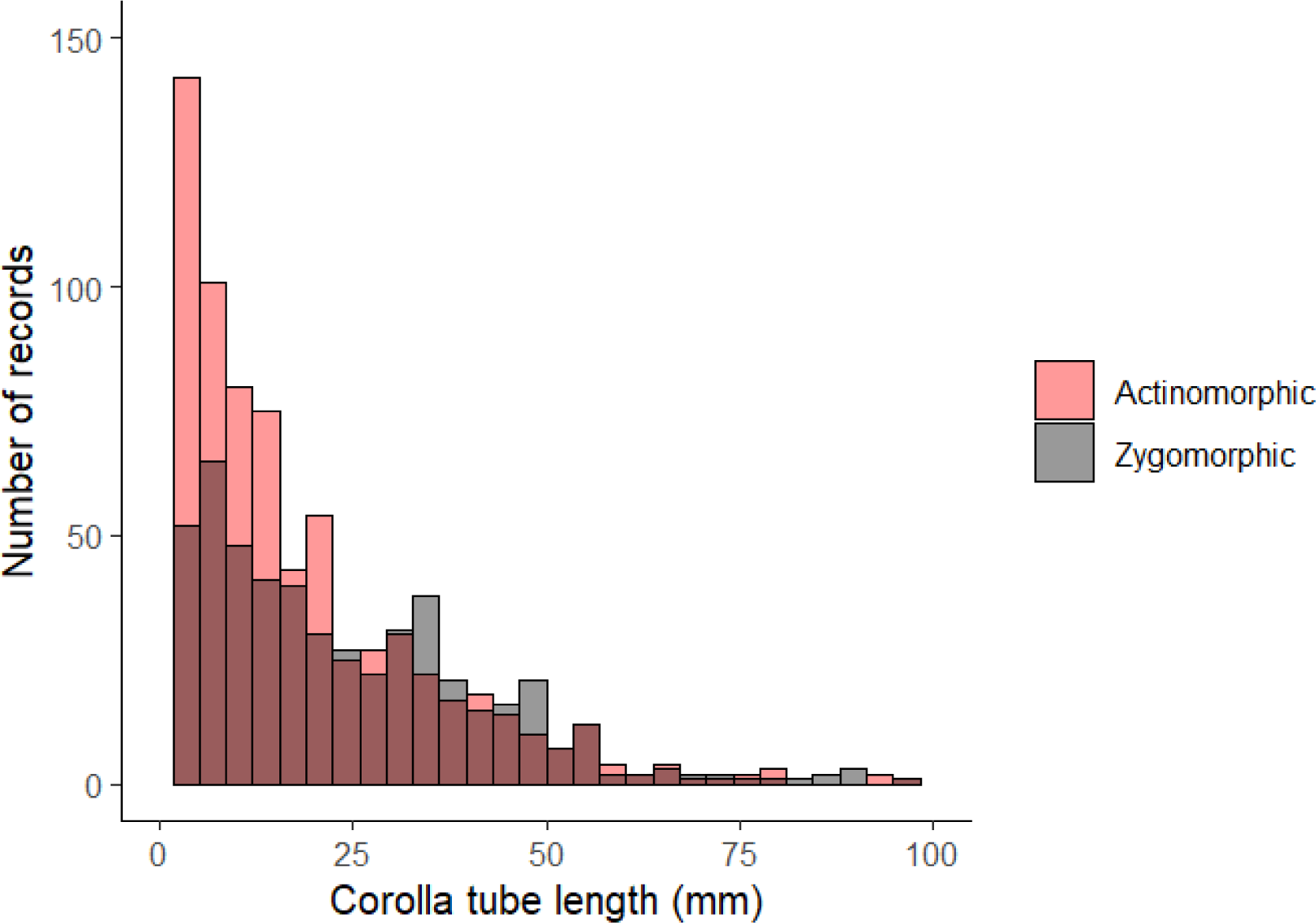
The distribution of mean per-species corolla tube lengths in the analyzed data set.

### Tube length and symmetry as reward predictors

Information about both nectar and pollen production was available for 87 species in the dataset. The amounts of the two types of rewards were not correlated (Pearson’s correlation coefficient: -0.04, df = 85, P = 0.69). Corolla tube length, zygomorphy, and their interaction positively and significantly influenced sugar production (Figs. 2-3): sugar production was higher in zygomorphic flowers than in actinomorphic flowers, and increased with the length of corolla tube (PGLMM: Mean ± SE effect of tube length – 0.050 ± 0.010, P< 0.001, R2_lik = 0.36; effect of symmetry type – 1.343 ± 0.481, P<0.001, R2_lik = 0.26; effect of interaction – 0.027 ± 0.014, P = 0.04).

**Figure 2:**
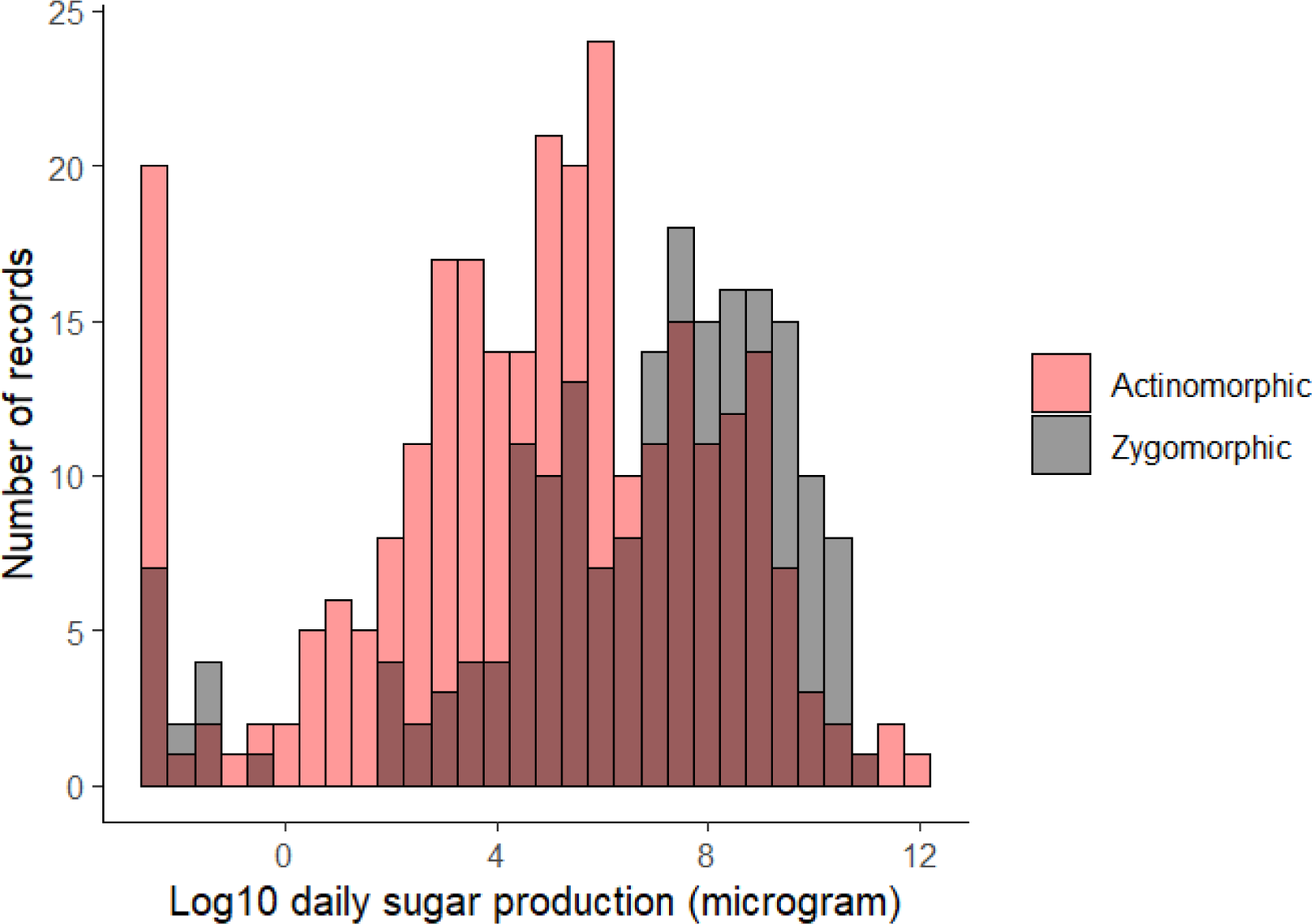
The distribution of daily sugar production rates. Negative values on the x-axis correspond to <1 microgram sugar/day.

**Figure 3:**
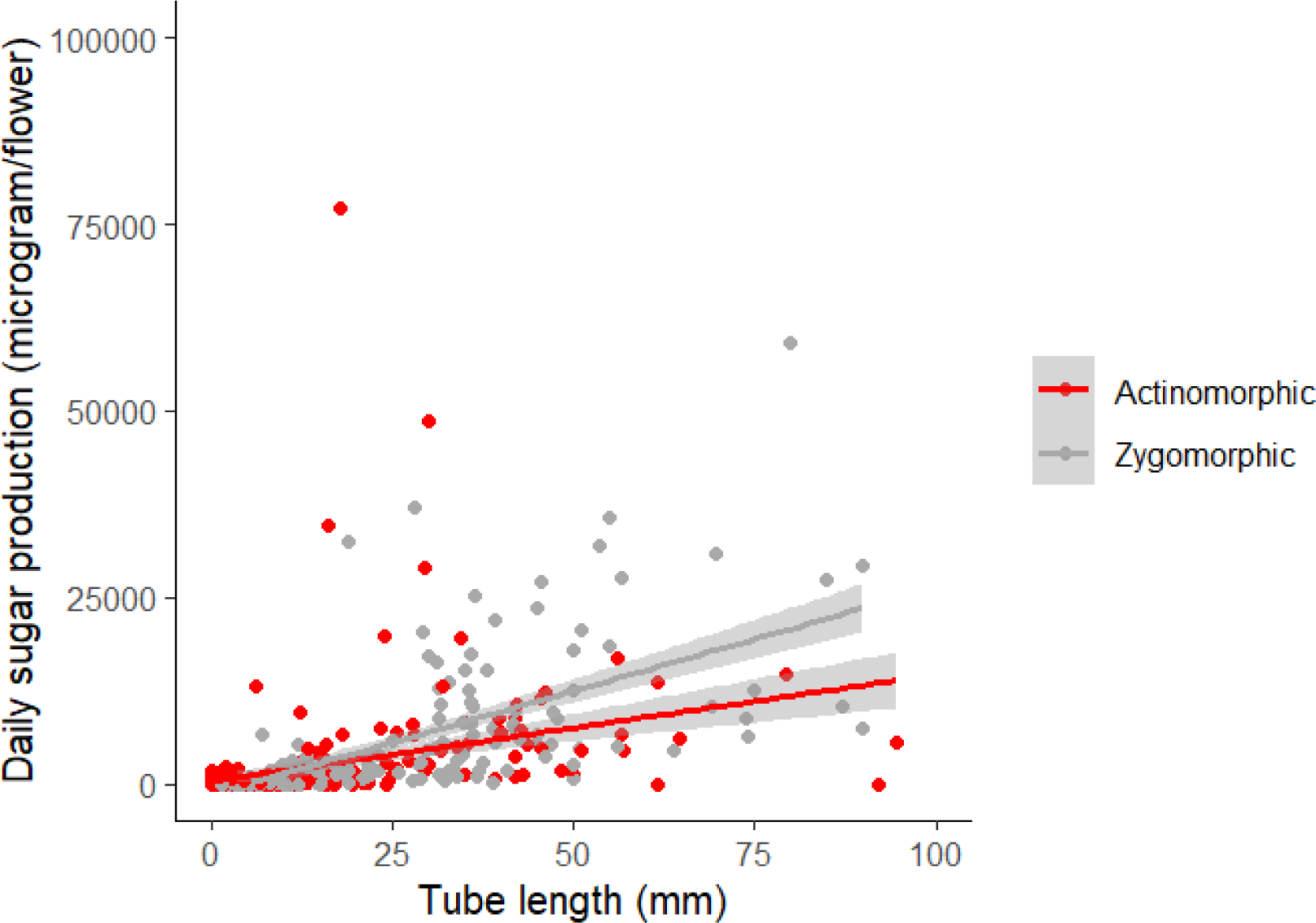
Daily sugar production rates vs. floral tube lengths.

Phylogeny, the random factor in the model, was as important as corolla tube length in the statistical model (R2_lik = 0.36). In contrast to their effect on sugar production, neither corolla tube length nor symmetry affected pollen counts (Figure 4. PGLMM: Mean ± SE effect of tube length – 0.005 ± 0.005, P = 0.371, R2_lik = 0.01; effect of symmetry type – -0.249 ± 0.153, P = 0.104, R2_lik = 0.08). Phylogeny was the most important predictor of pollen production (R2_lik = 0.43).

**Figure 4:**
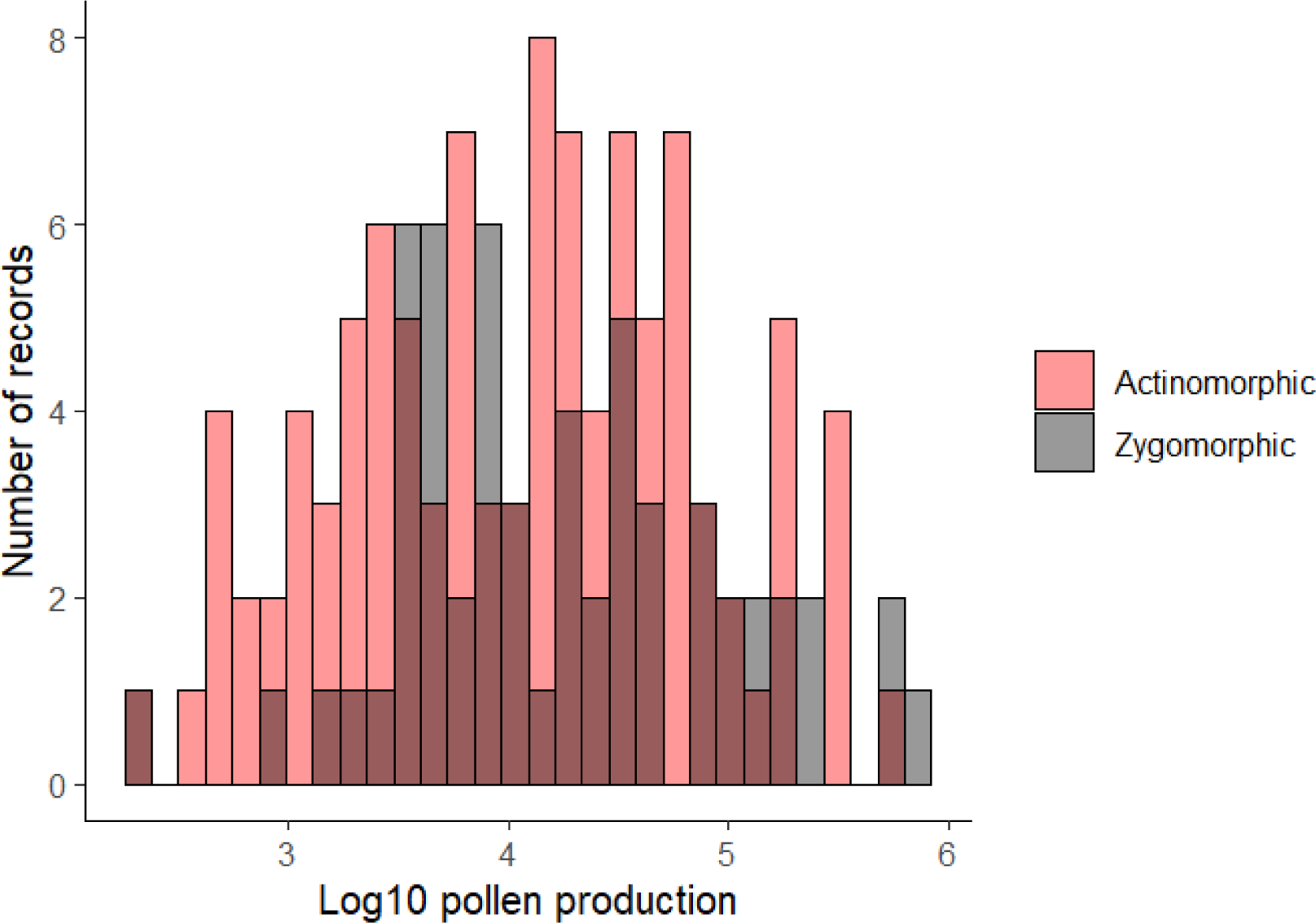
The distribution of per-flower pollen number

To better understand the role of phylogeny as reward predictor, we used generalized linear models to test the contribution of plant genus, nested within family, to the variation in sugar and pollen production. Both genus and family significantly (P < 0.001) affected sugar and pollen production. Genus had a larger effect than family on sugar production (partial eta-squared = 0.77 for genus, 0.58 for family), and a smaller effect than family on pollen production (partial eta-squared = 0.68 for genus, 0.76 for family). Mean ± SE per-family sugar and pollen production for the better-represented families (>20 species) in our dataset are provided in Table 1.

**Table 1:** Per-family mean (SE) per-flower daily sugar (microgram) and pollen (log no. pollen grains) production values.

| **Family** | **Nectar mean** | **Nectar SE** | **log(pollen) mean** | **log(pollen) SE** |
| --- | --- | --- | --- | --- |
| Aizoaceae |  |  | 5.540 | 0.114 |
| Apiaceae | 80.886 | 36.738 |  |  |
| Apocynaceae |  |  | 3.005 | 0.076 |
| Araceae |  |  | 4.016 | 0.161 |
| Asparagaceae | 11128.185 | 4674.410 |  |  |
| Asteraceae | 715.673 | 211.221 | 3.491 | 0.058 |
| Boraginaceae | 1584.893 | 917.551 |  |  |
| Brassicaceae | 109.426 | 32.025 | 4.213 | 0.076 |
| Bromeliaceae | 7327.772 | 1215.014 |  |  |
| Cactaceae |  |  | 5.298 | 0.126 |
| Campanulaceae | 2939.423 | 970.771 |  |  |
| Caprifoliaceae | 611.749 | 119.181 |  |  |
| Caryophyllaceae | 289.627 | 103.827 | 4.058 | 0.077 |
| Convolvulaceae |  |  | 3.307 | 0.074 |
| Ericaceae | 2123.128 | 593.280 | 4.953 | 0.130 |
| Fabaceae | 2019.320 | 487.275 | 4.269 | 0.047 |
| Gesneriaceae | 5073.087 | 727.156 |  |  |
| Lamiaceae | 793.868 | 100.865 | 3.717 | 0.072 |
| Malvaceae |  |  | 4.139 | 0.136 |
| Orchidaceae |  |  | 5.059 | 0.057 |
| Orobanchaceae |  |  | 4.716 | 0.071 |
| Phyllanthaceae |  |  | 3.673 | 0.089 |
| Plantaginaceae | 422.324 | 149.116 | 3.971 | 0.095 |
| Polemoniaceae |  |  | 3.525 | 0.061 |
| Ranunculaceae | 143.638 | 50.236 | 5.016 | 0.062 |
| Rosaceae | 154.230 | 44.230 | 5.010 | 0.074 |
| Rubiaceae | 3457.217 | 2217.246 | 3.722 | 0.110 |
| Solanaceae |  |  | 5.418 | 0.073 |

## DISCUSSION

Long corolla tubes and zygomorphy, two floral traits that increase pollination specialization, were associated with increased daily sugar production rates in our dataset. Inter-specific correlations between tube length and nectar production were previously found in smaller-scale analyses, only some of which controlled for phylogeny (Harder and Cruzan, 1990; Ornelas et al., 2007; Tavares et al., 2016; Martins and Johnson, 2013; Petanidou and Smets, 1995). In additional studies, corolla tube length also predicted nectar production in flowers within plant species (Harder and Cruzan, 1990) and even within individuals (Benitez-Vieyra et al., 2014). Our results regarding corolla tube lengths are thus expected and confirmatory. By contrast, our finding of a positive effect of zygomorphy on nectar production is novel. Further, the interaction between symmetry and tube length found in our study (Figure 3) indicates that nectar production increases faster with tube length in zygomorphic flowers than in actinomorphic ones. Our results differ from a previous study, which found no association between symmetry type and floral food rewards (Ortiz et al., 2021). The effect of symmetry on sugar production was smaller than the effect of tube length, as reflected in its lower R^2^ value. The larger sample size used in the present study (454 plant species, vs. 98 species analyzed by Ortiz et al., 2021) may have allowed better detection of this relatively weak link between symmetry type and sugar production.

Zygomorphic flowers tend to have longer corolla tubes than radial ones (Figure 1). We used PGLMM models to address this confound, testing for the effect of each variable separately. As visualized in Figure 3, zygomorphy correlated with higher nectar production while controlling for tube length (a proxy of flower size). This result is consistent with an *evolutionary* interpretation, which predicts stronger selection for nectar production in flowers that are prone to pollinator limitation (Martins and Johnson, 2013). It also sheds new light on the finding that zygomorphic flowers have fewer pollinators than actinomorphic ones (Yoder, 2020). This higher specialization of zygomorphic flowers may owe in part to their longer average tube lengths. Recently, Stephens et al. (2024) found that zygomorphy is also associated with increased flower longevity. This may provide zygomorphic flowers with an additional mechanism to ensure sufficient pollinator visits despite their higher specialization. The *physiological constraints* and *evolutionary processes* interpretations are both compatible with the positive effect of tube length on floral nectar production.

Neither tube length nor symmetry predicted pollen production across the plant species in our dataset. These results partially agree with a previous study, which found no effect of symmetry, but a negative effect of flower depth, on pollen production in mountain plant communities in China (Nepal et al., 2023). Cunha et al. (2022) reported an increase in pollen production with latitude, and interpreted this trend as an insurance strategy against variable pollinator abundances in temperate climates. Cunha et al. (2022) also found increased frequencies of zygomorphy with latitude, while Nepal et al. (2023) found that pollen number slightly increased with elevation. The lack of association between symmetry pattern and pollen production in the present study may be due to latitude or elevation effects, which we did not analyze. An additional potential explanation involves conflicting selection pressures on pollen production in specialized flowers: on the one hand, selection is expected to favor increased investment in pollen production to mitigate pollinator limitation. On the one hand, the pollinators’ greater flower specialization and constancy are expected to increase the efficiency of pollen transfer and reduce pollen wastage. This can decrease the plants’ benefit from producing pollen in large excess.

Floral visual display traits that correlate with nectar or pollen production are potentially honest signals of reward, regardless of the mechanism that underlies the correlation. Our analysis suggests that deep bilateral flowers can advertise nectar rewards, but not pollen rewards, to foragers. Controlled choice experiments can inform us whether insects indeed respond to these signals when foraging for nectar. With respect to flower symmetry, behavioral assays with different pollinators types show mixed responses, including preference for radially symmetrical flowers (Wignall et al., 2006) or no preference for either symmetry type (West and Lavery, 1998, Culbert and Forrest, 2016). Similar experiments are needed to assess the role of tube length as a visual signal. They should include choice assays with naïve foragers to test for innate preferences, and comparisons of pollinators’ success in associating either long or short corolla tubes with food rewards.

The morphological traits that we tested did not predict pollen number, hence cannot be considered as honest reward signals for pollen-foraging insects. Flower size, on the other hand, reliably predicts pollen rewards at the plant community level (Ortiz et al., 2021). Plant taxonomy, particularly family-level affiliation, seems to be an important predictor of pollen quantity (Harder and Johnson, 2023, this study) and chemical composition (Weiner et al., 2010). Whether and how pollen foragers use family-specific chemical or visual cues to select their food plants deserves further study.

## ACKNOWLEDGMENTS

The authors thank The Israel Science Foundation (grant No. 516/22 to TK) for financial support.

## DATA AVAILABILITY

Upon acceptance of the paper, the raw data used for the analyses will be posted at https://tamarkeasarlab.weebly.com/data-sets.html.

## Notes

### Competing Interest Statement

The authors have declared no competing interest.

